# Cohesin is involved in transcriptional repression of stage-specific genes in the human malaria parasite

**DOI:** 10.1101/2022.07.21.500927

**Authors:** Catarina Rosa, Parul Singh, Ameya Sinha, Peter R Preiser, Peter C Dedon, Sebastian Baumgarten, Artur Scherf, Jessica M Bryant

## Abstract

The most virulent human malaria parasite, *Plasmodium falciparum*, has a complex life cycle between its human host and mosquito vector. Each stage is driven by a specific transcriptional program, but with a relatively high ratio of genes to specific transcription factors, it is unclear how genes are activated or silenced at specific times. The *P. falciparum* genome is relatively euchromatic compared to the mammalian genome, except for specific genes that are uniquely heterochromatinized via HP1. There seems to be an association between gene activity and spatial organization; however, the molecular mechanisms behind genome organization are unclear. While *P. falciparum* lacks key genome-organizing proteins found in metazoans, it does have all core components of the cohesin complex. In other eukaryotes, cohesin is involved in sister chromatid cohesion, transcription, and genome organization. To investigate the role of cohesin in *P. falciparum*, we combined genome editing, mass spectrometry, chromatin immunoprecipitation and sequencing (ChIP-seq), and RNA sequencing to functionally characterize the cohesin subunit Structural Maintenance of Chromosomes protein 3 (SMC3). SMC3 knockdown in early stages of the intraerythrocytic developmental cycle (IDC) resulted in significant upregulation of a subset of genes involved in erythrocyte egress and invasion, which are normally expressed at later stages. ChIP-seq of SMC3 revealed that over the IDC, enrichment at the promoter regions of these genes inversely correlates with their expression and chromatin accessibility levels. These data suggest that SMC3 binding helps to repress specific genes until their appropriate time of expression, revealing a new mode of stage-specific, HP1-independent gene repression in *P. falciparum*.

## INTRODUCTION

The most virulent human malaria parasite, *P. falciparum*, has a significant impact on human health in endemic regions (World Health Organization, 2020). The approximately 48-hour intraerythrocytic developmental cycle (IDC) takes place in the human blood and is responsible for all clinical symptoms of malaria. During the IDC, each parasite replicates by schizogony, giving rise to up to 36 daughter cells that egress out of the red blood cell (RBC) and begin a new round of infection (Cowman et al., 2016). Underlying parasite development across the IDC is a highly coordinated gene expression program in which transcription of most genes peaks when the corresponding protein is required (Bozdech et al., 2003; Painter et al., 2018). Since the major limiting step for gene expression is transcription initiation (Caro et al., 2014), one possibility is that expression patterns result from a precisely timed production and/or binding of sequence-specific transcription factors (TFs). While recent studies of chromatin accessibility show evidence for dynamic exposure of potential transcription factor binding sites upstream of genes, the *P. falciparum* genome encodes few sequence-specific TFs compared to other eukaryotes, accounting for less than 1% of the protein-coding genes (Balaji et al., 2005; Campbell et al., 2010; Toenhake et al., 2018).

The majority of the *P. falciparum* genome is in a transcriptionally permissive, euchromatic state, with histone acetylation and deacetylation being the main predictors of gene activation or repression, respectively (Salcedo-Amaya et al., 2009; Trelle et al., 2009). Exceptions to this rule include multigene families encoding variant surface antigens, which are uniquely heterochromatinized via heterochromatin protein 1 (HP1) and form clusters at the nuclear periphery (Lopez-Rubio et al., 2009; Ralph et al., 2005). The recent application of genome-wide chromosome conformation capture techniques (Hi-C) confirmed close association of these multigene families, the clustering of centromeres and telomeres at opposite sides of the nucleus, and co-localization of active ribosomal DNA (rDNA) genes (Ay et al., 2014; Bunnik et al., 2019). In addition to the strong clustering of HP1-enriched multigene families and highly transcribed rDNA units, genes with similar expression profiles were also found to associate in a spatiotemporal manner during the IDC (Ay et al., 2014). Indeed, certain gene families appear to change their localization within the nucleus between the IDC (rings, trophozoites, and schizonts), the transmission from human to mosquito (early and late gametocytes), and from mosquito to human (sporozoites) (Bunnik et al., 2018).

Although this growing body of evidence shows that specific genes and genomic features associate at specific times in the *P. falciparum* life cycle, the factors responsible for this organization are largely unknown. Protein factors including actin and HP1 were shown to play a role in the organization and transcriptional regulation of the *var* multigene family (Lopez-Rubio et al., 2009; Ralph et al., 2005; Q. Zhang et al., 2011). More recently, an architectural factor, the high-mobility-group-box protein 1 (*Pf*HMGB1) was found to play a role in the nuclear organization of centromeres, and knockdown led to defects in *var* gene transcription (Lu et al., 2021). However, architectural factors linking chromosomal organization to the strict spatio-temporal transcriptional regulation of HP1-independent genes remain to be uncovered.

Although *P. falciparum* lacks lamins and CCCTC-binding factor (CTCF) – key genome organizing proteins in metazoans (Batsios et al., 2012; Heger et al., 2012) – it encodes the functionally uncharacterized putative orthologues of the core components of the cohesin complex: Structural Maintenance of Chromosomes protein 1 (SMC1, PF3D7_1130700), SMC3 (PF3D7_0414000), and an α-kleisin subunit (RAD21) (PF3D7_1440100) (Gardner et al., 2002). Among eukaryotes investigated, cohesin is a multiprotein complex that performs multiple different functions that primarily rely on its ability to topologically entrap strands of DNA (reviewed in (Dorsett & Ström, 2012; Perea-Resa et al., 2021; Uhlmann, 2016)). SMC1 and SMC3 each contain a hinge domain, which facilitates dimerization between the two proteins, and an ATPase head domain (Fig. 1A). Association with RAD21 at the SMC1/3 head domains results in a ring-like structure (Fig. 1A) that is able to both entrap DNA (Gligoris et al., 2014; Huis in ’t Veld et al., 2014) and extrude DNA loops (Davidson et al., 2019; Kim et al., 2019).

**Figure 1.**
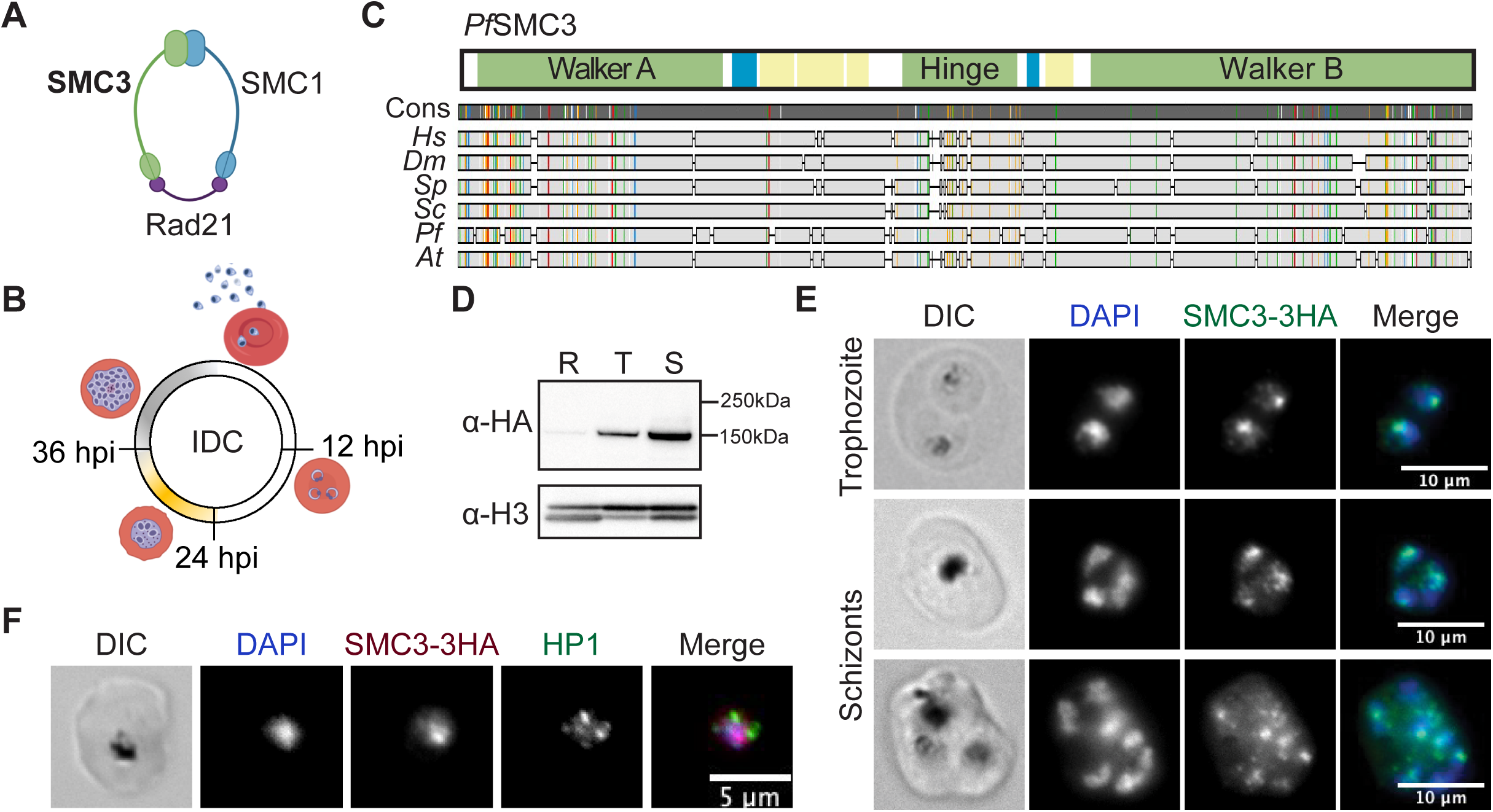
SMC3 is expressed across the IDC and localizes to HP1-independent nuclear foci. **(A)** Cohesin complex subunits annotated in *P. falciparum* (Gardner et al., 2002). Image prepared with BioRender.com. **(B)** Schematic of *P. falciparum* intraerythrocytic developmental cycle (IDC). Yellow, approximate timing of DNA replication; Grey, approximate duration of schizogony (modified from Ganter *et al*., 2017 and Matthews *et al*., 2018). Time points in this study – 12 hpi (ring), 24 hpi (trophozoite), and 36 hpi (schizont) – are indicated. **(C)** Alignment of *P. falciparum* (*Pf*) SMC3 (*Pf*SMC3) with SMC3 protein sequences in *H. sapiens* (*Hs*), *D. melanogaster* (*Dm*), *S. pombe* (*Sp)*, *S. cerevisiae* (*Sc*), and *A. thaliana* (*At*). A schematic of *Pf*SMC3 domain architecture is shown above. Coiled-coil domains are in yellow, low complexity regions are in blue, and other structured domains are annotated and in green. Sequence consensus (“Cons”) is indicated by the grey bar with colors representing regions of 100 % agreement between the aligned sequences. Image prepared with Geneious Prime 2020.0.3. **(D)** Western blot analysis of nuclear extracts of ring (R), trophozoite (T), and schizont (S) stages from a synchronous population of SMC3-3HA-*glmS* parasites. SMC3-3HA is detected with an anti-HA antibody. An antibody against histone H3 is used as a control for the nuclear extract. Molecular weights are shown to the right. The SMC3-3HA has a predicted molecular weight of 147.3 kDa (3.3 kDa corresponding to the 3HA tag). **(E)** and **(F)** Immunofluorescence assays of fixed RBCs infected with trophozoite or schizont stage SMC3-3HA-*glmS* parasites. DNA was stained with DAPI (blue) and SMC3-3HA was detected with anti-HA (green in E and magenta in F) antibody. HP1 was detected with anti-HP1 antibody (green in F). DIC, differential interference contrast. Scale bars equal 10 µm (E) and 5 µm (F).

The most well-characterized role of cohesin is in holding replicated sister chromatids together to ensure faithful chromosome segregation during cell division (Michaelis et al., 1997). Cohesin is loaded onto chromosomes during G_1_ or S phase, but in early mitosis, most of it is removed except for at centromeric and pericentromeric regions. This final pool is removed at the onset of anaphase to facilitate chromatid separation (reviewed in (Mirkovic & Oliveira, 2017; Peters & Nishiyama, 2012)). In the IDC of *P. falciparum*, asexual replication is accomplished through endocyclic schizogony, during which asynchronous rounds of DNA replication and mitosis lead to multinucleated cells, all in the absence of chromosome condensation (Fig. 1B). Schizogony culminates with a final round of nuclear division before cytokinesis (Klaus et al., 2022; Rudlaff et al., 2020). While much recent progress has been made in elucidating the mechanisms behind this unique cell division, many questions remain.

In contrast to mitosis, cohesin binding during G_1_ phase or in non-dividing cells was found to be more dynamic (Eichinger et al., 2013; Gerlich et al., 2006). In fact, in more recent years, the cohesin complex and the regulatory proteins that control the loading and unloading of the complex to DNA were found to play a role in shaping chromosomal architecture and thus, transcription, during interphase. In mammalian cells, cohesin and CTCF are often found at the boundaries of topologically associating domains (TADs) (Dixon et al., 2012; Nuebler et al., 2018; Rao et al., 2017; Wutz et al., 2017). A TAD is a region of the genome that preferentially interacts with itself in comparison with the rest of the genome (reviewed in (Dekker & Heard, 2015)). Importantly, TADs have emerged as functional structures involved in the regulation of cell type- and developmental stage-specific transcriptional programs, most likely via the correct pairing of enhancers with promoters (reviewed in (Dixon et al., 2016; Perea-Resa et al., 2021)). While the *P. falciparum* genome is not organized into TADs, as they are defined in metazoans, it does feature long-range inter- and intra-chromosomal that are involved in transcriptional control (Ay et al., 2014; Bunnik et al., 2019; Q. Zhang et al., 2011).

In *Plasmodium*, the physical association of SMC1, SMC3, and a protein containing the Rad21/Rec8-like N-terminal domain has been described (Hillier et al., 2019). Recently, a preliminary characterization of *Pf*SMC3 was carried out using an antibody generated in-house (Batugedara et al., 2020). Chromatin immunoprecipitation and sequencing (ChIP-seq) in trophozoites revealed that SMC3 was enriched at centromeric regions (Batugedara et al., 2020). In the present study, we use genome editing, mass spectrometry, and ChIP- and RNA-seq to functionally characterize the cohesin subunit SMC3 in interphase transcriptional regulation during the IDC. We show that while SMC3 is constantly present at centromeres across the IDC (Batugedara et al., 2020), it binds dynamically to the promoters of a specific subset of genes that are upregulated in its absence. Our findings represent a new mode of transcriptional repression in *P. falciparum*.

## RESULTS

### SMC3 is expressed across the IDC and localizes to HP1-independent nuclear foci

In the *P. falciparum* genome, three putative core cohesin subunits have been annotated: SMC1 (PF3D7_1130700), SMC3 (PF3D7_0414000), and a protein with the N-terminal Rad21/Rec8 domain (PF3D7_1440100) (Fig. 1A). A comparative sequence analysis showed that, of these three subunits, *Pf*SMC3 shares the highest sequence similarly and identity to its orthologues in *H. sapiens*, *D. melanogaster*, *S. cerevisiae*, *S. pombe,* and *A. thaliana* (Fig. 1C). A Pfam domain analysis (Mistry et al., 2021) showed an overall conserved domain architecture: an N-terminal Walker A motif-containing domain, a central hinge domain, and a C-terminal Walker B motif-containing domain (Fig. 1C). Given the conserved nature of *Pf*SMC3, we decided to investigate its function *in vivo*.

We used CRISPR/Cas9 genome editing (Ghorbal et al., 2014) to add a 3x hemagglutinin (3HA) epitope tag-encoding sequence followed by a *glmS* ribozyme-encoding sequence at the 3’ end of *smc3* (SMC3-3HA-*glmS*), which allows for inducible knockdown (Prommana et al., 2013). Immunoprecipitation followed by liquid chromatography-mass spectrometry (IP LC-MS/MS) of SMC3-3HA confirmed the interaction of SMC1, SMC3, and RAD21 previously reported in *P. falciparum* (Hillier et al., 2019; Batugedara et al., 2020) (Table 1). A Stromal Antigen (STAG) domain-containing protein (PF3D7_1456500) was also enriched in the SMC3-3HA IP LC-MS/MS, suggesting that a fourth cohesin subunit (STAG1/2 in *H. sapiens* and Scc3 in *S. cerevisiae*) is present in the *P. falciparum* cohesin complex (Table 1).

Western blot analysis of a synchronous bulk population of SMC3-3HA*-glmS* parasites showed that SMC3 is expressed across the IDC, but increases in abundance from ring to schizont stage (Fig. 1D). The presence of SMC3 in both ring and trophozoite stages suggests that cohesin is playing a role in interphase parasites (i.e. outside schizogony) and perhaps even before the onset of S phase, which is believed to take place after 24 hpi (Arnot et al., 2011; Ganter et al., 2017; Stanojcic et al., 2017) (Fig. 1B). Immunofluorescence assay (IFA) corroborated the nuclear localization, revealing a focus of SMC3-3HA at the nuclear periphery in trophozoite and schizont stages (Fig. 1E). While these foci are reminiscent of the heterochromatic *var* gene clusters at the nuclear periphery, no co-localization was observed between SMC3 and HP1 foci in trophozoite stage (Fig. 1F). It was not possible to detect SMC3 in ring stage and early trophozoite parasites with IFA, possibly due to the low abundance of the protein at this stage.

### SMC3 binds stably to centromeres, but dynamically to other genes across the IDC

To determine the genome-wide binding pattern of SMC3 across the IDC, ChIP-seq was performed in a synchronous clonal population of SMC3-3HA-*glmS* parasites at 12 (ring), 24 (trophozoite), and 36 (schizont) hours post invasion (hpi). Using the macs2 peak calling algorithm (Y. Zhang et al., 2008), we obtained 1,164, 1,614, and 1,027 significant peaks at 12, 24, and 36 hpi, respectively (Table 2). Most striking was the SMC3 enrichment at centromeric regions at all time points, a phenomenon that was previously reported for trophozoite stages (Batugedara et al., 2020) (Fig. 2 A, B). Comparison of the SMC3 peaks with the centromeric regions defined in (Hoeijmakers et al., 2012) revealed extensive overlap (Table 3). SMC3 peak enrichment in centromeric regions was significantly higher than that of the peaks associated with the rest of the genome at 12, 24, and 36 hpi (*P* < 0.0001). Interestingly, we observed a decrease in SMC3 enrichment at the centromeric regions from 24 to 36 hpi, a time that corresponds to the transition into mitosis (Fig. 2C, Table 3).

**Figure 2.**
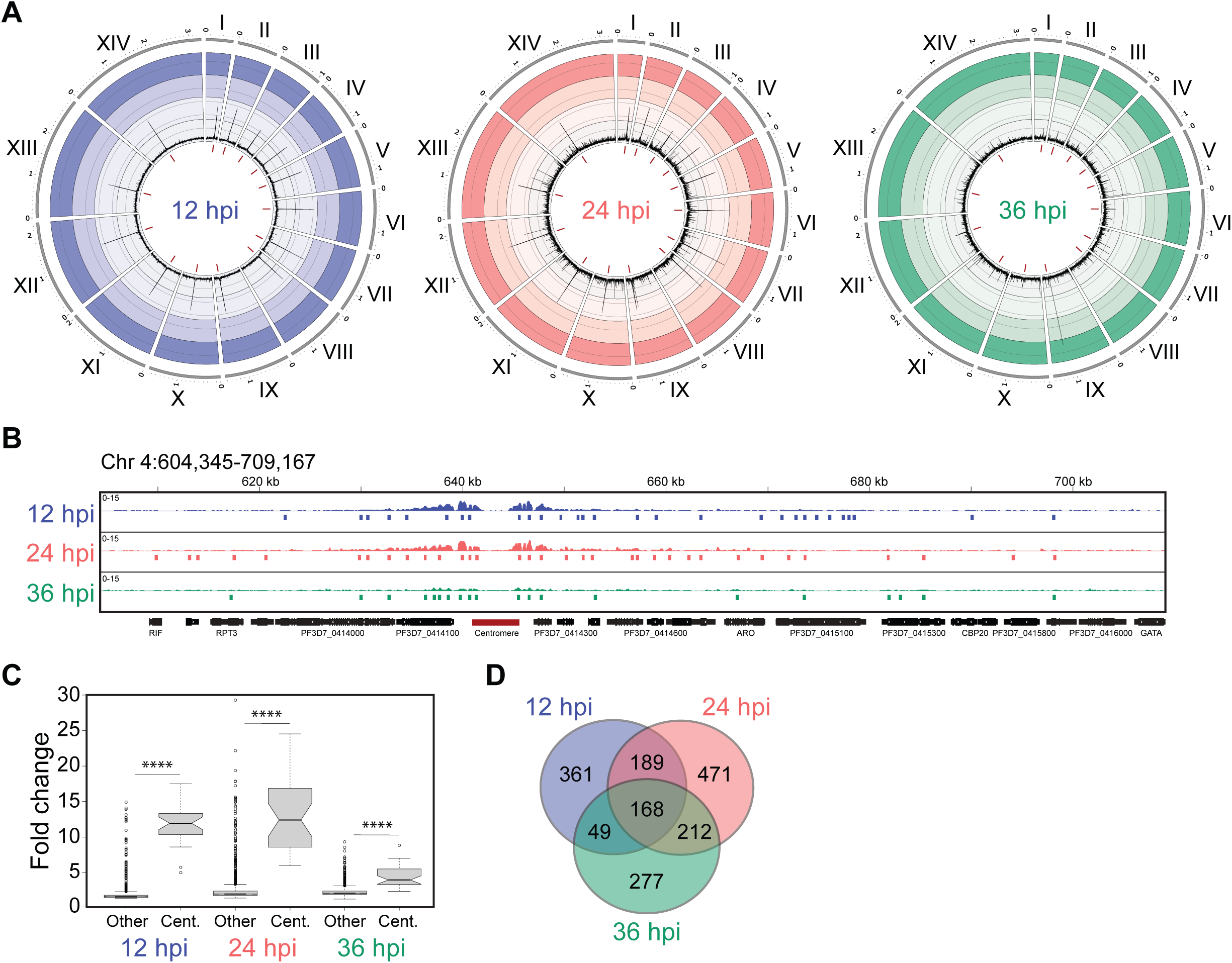
SMC3 binds stably to centromeres, but dynamically to other genes across the IDC. **(A)** Circos plot of ChIP-seq data showing genome-wide SMC3 binding across the IDC. For 12 (blue), 24 (coral), and 36 (green) hpi, the 14 chromosomes are represented circularly by the outer gray bars, with chromosome number indicated in roman numerals and chromosome distances (Mbp) indicated in Arabic numerals. Enrichment ratio (ChIP/input) is shown as average reads per million (RPM) over bins of 1,000 nucleotides. The maximum *y*-axis value is 24. Centromeric regions are represented by red bars in the innermost circle. **(B)** Zoomed-in view of ChIP-seq data corresponding to chromosome 4 (604,345 - 709,167 bp), including the centromere (represented with dark red line below the *x*-axis). For 12 (blue), 24 (coral), and 36 (green) hpi the *y*-axis is enrichment (ChIP/Input), with vertical lines below representing significant peaks obtained from peak calling algorithm macs2 (*q*-value < 0.05). The *x*-axis is DNA sequence, with genes represented by black boxes indented to delineate introns and labeled with white arrowheads to indicate transcription direction. **(C)** Box plot comparing the distribution of peak enrichment (fold change, ChIP/Input) between centromeric (Cent.) regions and extra-centromeric (Other) regions of the genome for 12, 24, and 36 hpi. Peaks were called with macs2 (*q*-value < 0.05). Center line, median; box limits, first and third quartiles; whiskers, 1.5× interquartile range. Wilcoxon test was used for statistical analysis. **** = adjusted *P*-value < 0.0001. **(D)** Venn diagram showing overlap between SMC3 peak-associated genes at 12 (blue), 24 (coral), and 36 (green) hpi. Closest unique protein coding genes to the extended SMC3-3HA peak summit (+/- 500 bp) at 12, 24, and 36 hpi are shown in Table 4.

While quantification of the SMC3 peaks showed the largest enrichment in the centromeric and pericentromeric regions, there were significant SMC3 peaks across other genomic locations at all time points (Table 2). SMC3 peaks were found in intergenic and intragenic regions closest to 767, 1,044, and 708 protein coding genes at 12, 24, and 36 hpi, respectively (Table 4). Of all genes within ±500 base pairs (bp) of an SMC3 peak, 168 were bound by SMC3 across all three time points (Fig. 2D). However, most SMC3-bound genes showed a dynamic binding pattern, with a peak present at only one or two time points (Fig. 2B,D). Gene ontology (GO) enrichment analysis showed that genes associated with SMC3 peaks at 12 hpi were not significantly represented by a specific GO term category (Table 5). However, genes associated with SMC3 peaks at 24 and 36 hpi were most significantly represented by biological process categories such as “obsolete pathogenesis” (*q* = 1.2 x 10^-19^ and 3.3 x 10^-21^, respectively), “cell-cell adhesion” (*q* = 1.2 x 10^-19^ and 4.7 x 10^-23^, respectively), “response to host” (*q* = 1.21 x 10^-11^ and 1.43 x 10^-13^, respectively), and “antigenic variation” (*q* = 8.1 x 10^-12^ and 2.7 x 10^-13^, respectively) (Table 5). These categories include many genes in common such as *var* and *rif* genes, which encode proteins that are exported to the surface of the host red blood cell to facilitate adhesion to the host microvasculature (reviewed in (Scherf et al., 2008)). Genes associated with SMC3 peaks at 24 hpi were also significantly represented by the biological process categories “entry into host” (*q* = 0.014) and “exit from host” (*q* = 0.031). These categories include genes that are involved in invasion of or egress from the red blood cell such as *ralp1* (PF3D7_0722200) (Haase et al., 2008), *rhoph3* (PF3D7_0905400) (Sherling et al., 2017), and *msp1* (PF3D7_0930300) (O’Donnell et al., 2000, 2001).

While peak calling analysis is informative, the diverse functional categories of genes associated with SMC3 peaks makes it difficult to determine if SMC3 plays a specific role in transcriptional regulation or binds randomly throughout genic regions to facilitate a role in mitosis-related chromosome organization. Thus, functional analysis was required to elucidate a potential transcriptional function for SMC3 binding.

### SMC3 inducible knockdown results in deregulation of genes during interphase

To gain insight into the role of SMC3 during interphase, we performed an inducible knockdown of SMC3 using the *glmS* ribozyme system (Prommana et al., 2013). An SMC3-3HA-*glmS* clone was tightly synchronized and split, and glucosamine was added to one half for 96 hours (2 cell cycles), as knockdown at the protein level could not be achieved after a single cell cycle (Supp. Fig. 1). Simultaneously, a wild-type (WT) clone from the parent 3D7 strain was synchronized and treated in the same way to account for transcriptional changes due to the presence of glucosamine. After another round of synchronization, parasites were harvested at 12 and 24 hpi, and western blot analysis revealed an SMC3-3HA knockdown at the protein level in nuclear extracts at both time points (Fig. 3A).

**Figure 3.**
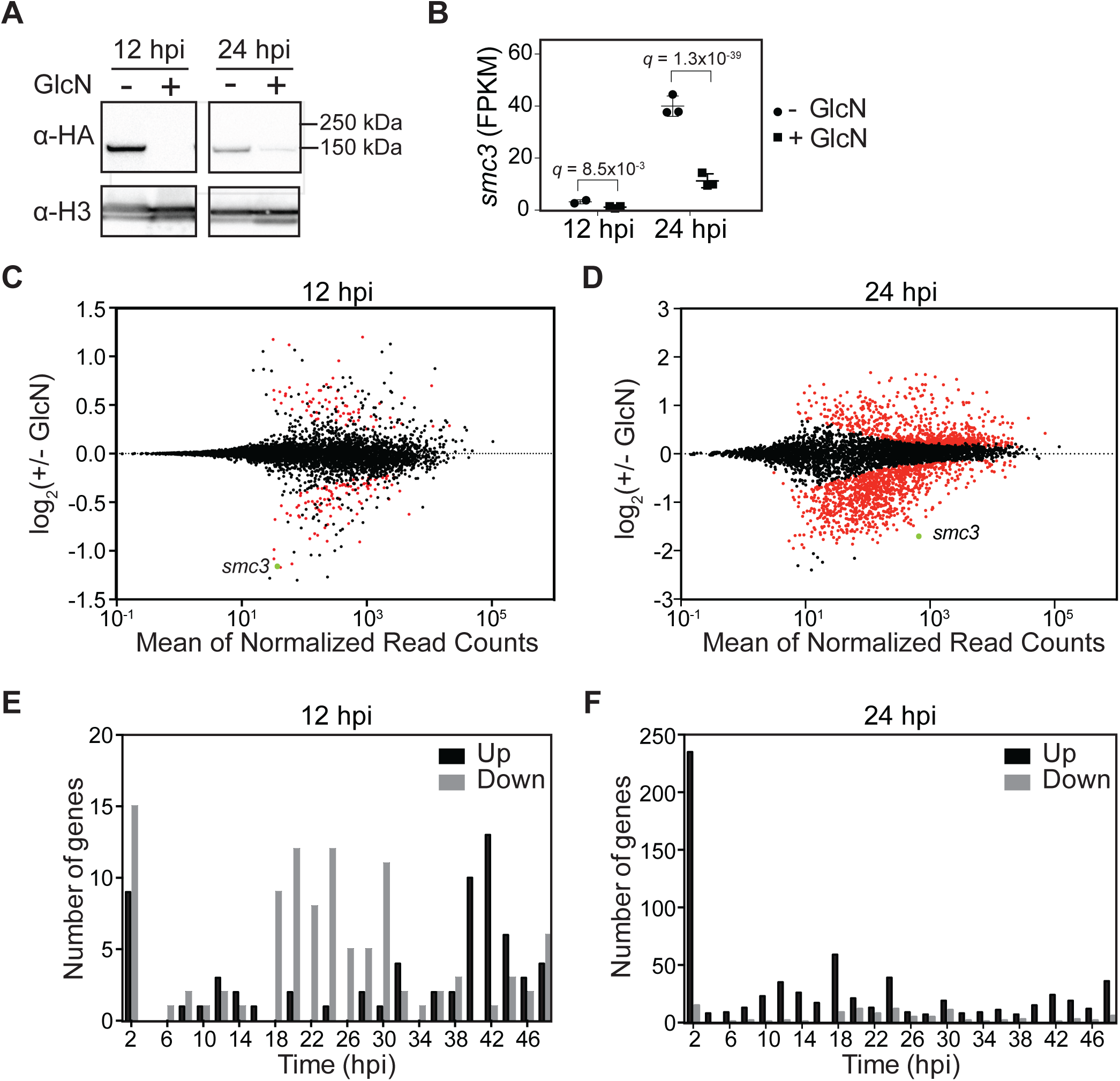
SMC3 inducible knockdown results in deregulation of genes during interphase. **(A)** Western blot analysis of nuclear extracts at 12 and 24 hpi from a clonal population of SMC3-3HA-*glmS* parasites in the absence (-) or presence (+) of glucosamine (GlcN). SMC3-3HA is detected with an anti-HA antibody. An antibody against histone H3 is used as a control for the nuclear extract. Molecular weights are shown to the right. **(B)** RNA-seq of an SMC3-3HA-*glmS* clone shows *smc3* transcript levels (FPKM) at 12 (*q* = 8.5 x 10^-3^) and 24 (*q* = 1.3 x 10^-39^) hpi in the absence (circles) or presence (squares) of glucosamine (GlcN). *P*-values are calculated with a Wald test for significance of coefficients in a negative binomial generalized linear model as implemented in DESeq2 (Love et al., 2014). *q* = Bonferroni corrected *P*-value. Corresponding data can be found in Tables 6 and 7. **(C)** and **(D)** MA plots of log_2_(glucosamine-treated/untreated, M) plotted over the mean abundance of each gene (A) at 12 hpi **(C)** and 24 hpi **(D)**. Transcripts that were significantly higher (above *x*-axis) or lower (below *x*-axis) in abundance in the presence of glucosamine are highlighted in red (*q* ≤ 0.1). *smc3* is highlighted in green. Three replicates were used for untreated and glucosamine-treated parasites, with the exception of the untreated 12 hpi parasites, for which there were two replicates. *P-*values were calculated with a Wald test for significance of coefficients in a negative binomial generalized linear model as implemented in DESeq2 (Love et al., 2014). *q = B*onferroni corrected *P-*value. **(E)** and **(F)** Frequency plots showing the time in the IDC (hpi) of peak transcript level (comparison to transcriptomics time course in (Painter et al., 2018)) for genes that are significantly downregulated (grey) or upregulated (black) following SMC3 knockdown at 12 hpi **(E)** and 24 **(F)** hpi.

We then performed RNA-seq followed by differential expression analysis for the untreated and glucosamine-treated SMC3-3HA-*glmS* and WT parasites, which confirmed a significant knockdown of SMC3 at the transcript level in the SMC3-3HA-*glmS* parasites: 55% (*q* = 8.5 x 10^-3^) at 12 hpi and 69% (*q* = 1.3 x 10^-39^) at 24 hpi (Tables 6 and 7, Fig. 3B). To remove potential artifacts of glucosamine treatment, genes that were significantly up- or downregulated in the glucosamine-treated WT parasites at 12 and 24 hpi (Tables 8 and 9) were filtered out of the datasets for significantly up- and downregulated genes in the SMC-3HA-*glmS* parasites at the corresponding time points (Supp. Fig. 2). After this filtering, 104 and 932 genes were significantly downregulated at 12 and 24 hpi, respectively (Tables 10 and 11, Fig. 3 C,D), and 67 and 674 genes were significantly upregulated at 12 and 24 hpi, respectively (Tables 10 and 11, Fig. 3 C,D) in SMC3-3HA-*glmS* parasites. Comparison of our RNA-seq data to time course microarray data from (Bozdech et al., 2003), as in (Lemieux et al., 2009), showed that data from the untreated and glucosamine-treated parasites harvested at 12 hpi corresponded statistically to 12 hpi (Supp. Fig. 3). The untreated and glucosamine-treated parasites harvested at 24 hpi correspond statistically to approximately 18-19 hpi (Supp. Fig. 3). However, the glucosamine-treated parasites were slightly more advanced in the cell cycle than the untreated parasites at 24 hpi, which could account for the higher number of genes that were significantly differentially expressed at this time point.

To gain insight into the transcriptional function of SMC3, we performed a GO enrichment analysis of genes that were up- and downregulated specifically in response to SMC3 knockdown at 12 and 24 hpi. At 12 hpi, downregulated genes were most significantly represented by the biological process category of “protein insertion into membrane” (*q* = 0.017, Table 12), whereas at 24 hpi downregulated genes were most significantly represented by the categories of “chromosome organization” (*q* = 1.0 x 10^-3^, Table 13) and “chromosome segregation” (*q* = 1.0 x 10^-3^, Table 13).

For both time points, upregulated genes were most significantly represented by the biological process categories of “movement in host environment” (12 hpi: *q* = 1.8 x 10^-7^, Table 12; 24 hpi: *q* = 1.3 x 10^-5^, Table 13) and “entry into host” (12 hpi: *q* = 1.8 x 10^-7^, Table 12; 24 hpi: *q* = 1.3 x 10^-5^, Table 13). Genes included in these categories are involved in egress and invasion of the red blood cell (reviewed in (Cowman et al., 2012, 2017)). Indeed, a substantial percentage of invasion-related genes defined in (Hu et al., 2010) were significantly upregulated upon SMC3 depletion at 12 and 24 hpi (Table 14). Comparison of our RNA-seq data to the time course transcriptomics data from (Painter et al., 2018) revealed that SMC3 depletion at 12 hpi caused downregulation of genes that normally reach their peak expression in the trophozoite stage (18-30 hpi), with the majority of upregulated genes normally reaching their peak expression in the schizont and very early ring stages (40-2 hpi) (Fig. 3E). At 24 hpi, a similar trend is observed, with most downregulated genes normally peaking in expression in trophozoite stage (24-32 hpi) and the majority of upregulated genes peaking in expression at very early ring stage (2 hpi) (Fig. 3F).

### SMC3 is involved in transcriptional regulation of genes involved in invasion

To provide evidence for a direct function of SMC3 in the transcriptional regulation of these up- and downregulated genes, we compared our SMC3 ChIP-seq data to our RNA-seq data at 12 hpi. Metagene analysis from the ChIP-seq data showed that SMC3 was absent from the promoter regions of genes that are downregulated in response to its knockdown (Fig. 4A). In contrast, SMC3 was enriched in the promoter regions of genes that are upregulated in response to its knockdown (Fig. 4A). Indeed, this enrichment of SMC3 at the promoters of upregulated genes was present at 12 and 24 hpi, but not 36 hpi (Fig. 4B). Our data suggest that SMC3 binding has a direct effect on the transcription of genes that are upregulated in its absence, whether naturally or via knockdown.

**Figure 4.**
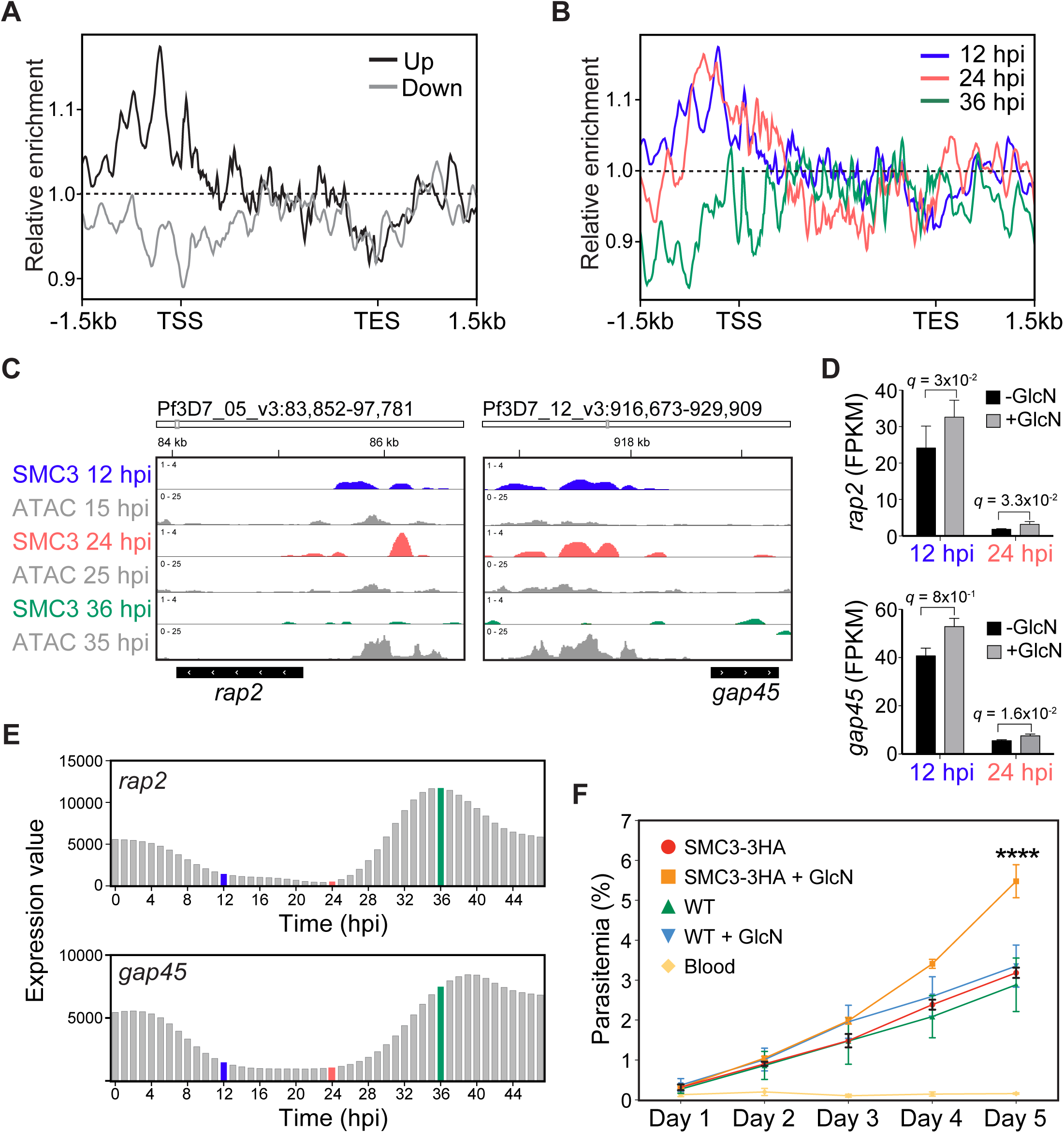
SMC3 is involved in transcriptional regulation of genes involved in invasion. Metagene plot showing average SMC3 enrichment (*y*-axis = ChIP/Input) in clonal SMC3-3HA-*glmS* parasites at 12 hpi from 1.5 kb upstream of the transcription start site (TSS) to 1.5 kb downstream of the transcription end site (TES) for genes that are significantly down- (grey) or upregulated (black) upon SMC3 knockdown. One replicate was used for the SMC3 ChIP-seq. **(A)** Metagene plot showing average SMC3 enrichment (*y*-axis = ChIP/Input) in clonal SMC3-3HA-*glmS* parasites at 12 (blue), 24 (coral), and 36 hpi (green) from 1.5 kb upstream of the transcription start site (TSS) to 1.5 kb downstream of the transcription end site (TES) for genes that are significantly upregulated upon SMC3 knockdown at 12 hpi. One replicate was used for the SMC3 ChIP-seq. **(B)** ChIP-seq data showing enrichment of SMC3 (ChIP/Input) at 12 (blue), 24 (coral), and 36 (green) hpi in clonal SMC3-3HA-*glmS* parasites at the *rhoptry-associated protein 2* (*rap2*, PF3D7_0501600) and the *glideosome-associated protein 45* (*gap45*, PF3D7_1222700) gene loci. The *x*-axis is DNA sequence, with the gene represented by a black box with white arrowheads to indicate transcription direction. One replicate was used for ChIP-seq. ATAC-seq data from closely corresponding time points (15, 25, and 35 hpi) from (Toenhake et al., 2018) are shown in grey, with the *y*-axis representing ATAC-seq (RPM)/gDNA (RPM). **(C)** RNA-seq of an SMC3-3HA-*glmS* clone shows transcript levels (FPKM) for *rap2* (PF3D7_0501600) at 12 (*q* = 3 x 10^-2^) and 24 (*q* = 3.3 x 10^-2^) hpi and *gap45* (PF3D7_1222700) at 12 (*q* = 8 x 10^-1^) and 24 (*q* = 1.6 x 10^-2^) hpi in the absence (black) or presence (grey) of glucosamine (GlcN). *P*-values are calculated with a Wald test for significance of coefficients in a negative binomial generalized linear model as implemented in DESeq2 (Love et al., 2014). *q* = Bonferroni corrected *P*-value. Corresponding data can be found in Tables 6 and 7. **(D)** Expression values of *rap2* (PF3D7_0501600) and *gap45* (PF3D7_1222700) genes across the IDC (indicated on the *x*-axis by hpi) from the transcriptomics time course in (Painter et al., 2018). Data corresponding to 12 (blue), 24 (coral), and 36 (green) hpi time points are highlighted. **(E)** Growth curve over five days of clonal SMC3-3HA-*glmS* and WT parasites in the absence or presence of glucosamine (GlcN). Glucosamine treatment was started 96 h (two cycles) before Day 1 to ensure SMC3 knockdown during the days sampled (Supp. Fig. 1). Uninfected red blood cells (Blood) served as reference of background. Error bars indicate standard deviation of three technical replicates in blood from two different donors (*n* = 6). A two-way ANOVA with Tukey post hoc test was used for statistical analysis. **** = adjusted *P*-value < 0.0001.

Because genes that are significantly upregulated upon SMC3 knockdown normally reach peak expression late in the cell cycle (Fig. 3E), are depleted of SMC3 at 36 hpi (Fig. 4B), and are most significantly represented by GO terms pertaining to invasion and egress (Tables 12,13), we hypothesized that SMC3 helps to repress these genes until their appropriate time of expression late in the cell cycle. Examples include the rhoptry-associated protein 2 (*rap2*, PF3D7_0501600) and glideosome-associated protein 45 (*gap45*, PF3D7_1222700). These genes show SMC3 enrichment at their promoter regions at 12 and 24 hpi, but not at 36 hpi (Fig. 4C), and depletion of SMC3 resulted in upregulation at both 12 and 24 hpi (Fig. 4D). Comparison of the SMC3 ChIP-seq data with published Assay for Transposase-Accessible Chromatin using sequencing (ATAC-seq) data (Toenhake et al., 2018) and mRNA dynamics data (Painter et al., 2018) from similar time points in the IDC revealed that SMC3 binding at the promoter regions of these genes inversely correlates with chromatin accessibility (Fig. 4C) and their mRNA levels (Fig. 4E), which both peak in schizont stages. These data are consistent with a role of SMC3 in repressing this gene subset until their appropriate time of expression in the IDC.

We hypothesized that the upregulation of invasion-related genes upon SMC3 knockdown might result in higher rates of invasion. Curiously, a comparison of growth between untreated and glucosamine-treated WT and SMC3-3HA-*glmS* parasites revealed a significantly higher growth rate in glucosamine-treated SMC3-3HA-*glmS* parasites in comparison to non-treated SMC3-3HA-*glmS*, treated 3D7 WT, and non-treated 3D7 WT parasites (*q* < 0.0001) (Fig. 4F). These data suggest that SMC3 knockdown results in a faster progression through the cell cycle or a higher rate of egress/invasion.

## DISCUSSION

Genome organization is key to transcriptional control and genome integrity. The human malaria parasite *P. falciparum* executes complex transcriptional programs and has a sophisticated genome organization considering that it encodes relatively few specific transcription factors and lacks key canonical genome organizing factors such as CTCF and lamins (Batsios et al., 2012; Heger et al., 2012; Ay et al., 2014; Bunnik et al., 2019). To investigate potential links between transcription and genome organization in this parasite, we have characterized SMC3, a key and conserved subunit of the multi-protein ring-shaped complex cohesin. In the organisms studied so far, cohesin plays diverse roles in genome organization such as sister chromatid cohesion during mitosis, transcription, and DNA damage repair (reviewed in (Perea-Resa et al., 2021)). Here, we used genome-wide approaches to elucidate the function of SMC3 in interphase transcription during the IDC of *P. falciparum*.

ChIP-seq over the course of the IDC revealed that SMC3 is most enriched in centromeric regions (Fig. 2A-C). In other eukaryotes, cohesin is also mostly enriched around the centromeres relative to the chromosome arms (Holzmann et al., 2019; Tanaka et al., 1999; Tomonaga et al., 2000). A reduction in centromere binding in late-stage parasites (Fig. 2A-C) might be due to the need for cohesin removal during the separation of sister chromatids, as has been observed in model eukaryotes (Uhlmann et al., 1999; Waizenegger et al., 2000). While *Plasmodium* does have a clear anaphase during which sister chromatids separate (Gerald et al., 2011), asynchronous mitosis may lead to a population of parasites in which cohesin is present or absent from centromeres to facilitate sister chromatid cohesion or separation, respectively. Our observation that SMC3 depletion does not inhibit parasite growth agrees with reports in *S. cerevisiae* and *D. melanogaster* in which normal growth and sister chromatid cohesion were achieved despite an 87% and 80% decrease in Rad21, respectively (Carvalhal et al., 2018; Heidinger-Pauli et al., 2010). These studies and ours suggest that only a small fraction of cohesin is needed to successfully complete mitosis.

Centromeric clustering in interphase nuclei has been observed in several eukaryotes including *S. cerevisiae, D. melanogaster*, and *H. sapiens* (reviewed in (Muller et al., 2019)). The functional importance of this spatial arrangement remains poorly understood; however, it has been shown that centromeric clustering is a relevant topological constraint that can affect transcription by preventing intrachromosomal arm interactions (Tolhuis et al., 2011). Studies in *P. falciparum* have demonstrated centromere clustering before and during schizogony, suggesting that this organization is needed during interphase and mitosis (Ay et al., 2014; Hoeijmakers et al., 2012). One architectural factor, *Pf*HMGB1, was recently shown to play a direct role in centromere organization in the nucleus (Lu et al., 2021). Although *Pf*HMGB1 binds predominantly to centromeres, its depletion led to the de-regulation of many different genes to which it was not bound, suggesting that global genome organization is important for transcriptional control at the local chromatin level (Lu et al., 2021). *Pf*HMGB1 knockout did not lead to blood stage parasite growth inhibition, indicating that other proteins, such as cohesin or *Pf*CenH3, play a role in centromere organization and mitosis.

In addition to its potential centromeric role, we discovered that SMC3 plays a direct, extra-centromeric role in the transcriptional control of specific genes during interphase. SMC3 bound dynamically at extra-centromeric genomic locations over the course of the IDC (Fig. 2, Table 2). We observed stage-specific SMC3 binding across the genome, including at genes that were then upregulated upon SMC3 depletion during interphase (Fig. 2B,D, Fig. 4A-D, Tables 2, 10, 11). In contrast, genes that were downregulated upon SMC3 depletion were not enriched for SMC3, suggesting an indirect effect (Fig. 4A). And while SMC3 peak-associated genes were significantly represented by GO terms related to antigenic variation at 24 and 36 hpi, significantly upregulated genes at 24 hpi did not show significant *q*-values for these or related GO terms. Upregulated genes are generally most highly expressed in late-stage parasites (Fig. 3E, Fig. 4E), a time when we observed natural depletion of SMC3 at their promoters (Fig. 4B,C). Importantly, while we observed a decrease in SMC3 binding at centromeric and pericentromeric regions in late-stage parasites, this was not a general trend across all SMC3 binding sites (Fig. 2C). These data suggest that SMC3 is specifically recruited to and evicted from specific subsets of genes to facilitate their repression and transcription, respectively, over the course of the IDC.

Genes that were significantly upregulated upon SMC3 depletion during interphase were enriched for GO terms related to egress from and invasion of the RBC (Tables 12 and 13). Indeed, out of a list of 63 invasion-related genes (Hu et al., 2010), 50% were among the genes that were upregulated upon SMC3 depletion during interphase (Table 14). Curiously, parasites depleted of SMC3 showed an increase in growth rate (Fig. 4F). It is possible that this phenotype is related to a potential role for SMC3 in mitosis or DNA repair, such as an as-yet unknown cell cycle checkpoint. However, this growth phenotype might also be the result of the upregulation of specific genes that allow for more efficient egress and invasion of new RBCs.

The mechanism by which SMC3 could repress specific genes in a stage-specific manner is unclear. In the context of interaction with CTCF, cohesin has been shown to impact transcription in opposite ways, by either preventing enhancer-promoter interactions (Nativio et al., 2009; Wendt et al., 2008) or by mediating specific enhancer-promoter loops (Kubo et al., 2021; Oh et al., 2021). In *P. falciparum,* the current genome-wide chromosome conformation capture (Hi-C) datasets do not provide evidence of typical enhancer-promoter interactions found in other eukaryotes (Ay et al., 2014; Bunnik et al., 2019). Moreover, in *S. cerevisiae,* a cohesin mutant resulted in the de-repression of genes located in subtelomeric regions, perhaps via disruption of local chromatin structure (Kothiwal & Laloraya, 2019). However, the invasion-related genes affected by *Pf*SMC3 are scattered across the genome (Table 14). One possibility is that cohesin binding to the promoter of a gene merely inhibits the transcriptional machinery from assembling. In light of the ability of cohesin to entrap multiple DNA molecules, another intriguing possibility is that it tethers invasion-related genes together in a cluster that renders their promoters inaccessible to specific activating factors until the appropriate time of transcription. Future high-resolution chromosome conformation capture studies will reveal a potential link between spatial association and transcriptional regulation of these SMC3-controlled genes.

It is also unclear how *Pf*SMC3 achieves binding specificity and how it is evicted from binding sites at specific times in the IDC. In other organisms studied, cohesin appears to need a DNA-binding factor to achieve sequence-specific binding (Kagey et al., 2010; Sasca et al., 2019; Wendt et al., 2008). A search for a specific binding motif within the promoter sequences of invasion-related genes bound by *Pf*SMC3 yielded no results, indicating that *Pf*SMC3 may associate with multiple factors to achieve specific binding. In *P. falciparum*, the AP2-I transcription factor (PF3D7_1007700) is involved in transcription of invasion-related genes via binding to the GTGCA motif, likely by interaction with the bromodomain protein 1 (*Pf*BDP1, PF3D7_1033700) (Santos et al., 2017). This complex could evict SMC3 from gene promoters or simply bind in its absence. Importantly, neither *ap2-I* nor *bdp1* are bound by SMC3 or are upregulated upon its depletion, suggesting that the observed upregulation of invasion-related genes upon SMC3 depletion in early-stage parasites is a direct effect. In addition, SMC3 depletion resulted in the upregulation of AP2-I-independent invasion-related genes such as RONs, EBLs and Rhs, which have an ACAACT motif in their promoter regions (Young et al., 2008) and may be activated by an as-yet unidentified transcription factor. Future studies will reveal the molecular machinery that regulates the stage-specific binding of cohesin.

The present study offers insight into the role of cohesin in the temporal regulation of genes in *P. falciparum*. While the role of H3K9me3/HP1 has been well established in the transcriptional repression of clonally variant gene families and *ap2-g*, this study identifies a new factor – SMC3 – involved in the repression of HP1-independent, stage-specific genes. Given the architectural nature of cohesin, this research provides a potential link between genome organization and transcriptional control in *P. falciparum*.

## MATERIALS AND METHODS

### Parasite culture

Blood stage 3D7 *P. falciparum* parasites were cultured as previously described in (Lopez-Rubio et al., 2009). Briefly, parasites were cultured in human RBCs supplemented with 10% v/v Albumax I (Thermo Fisher 11020), hypoxanthine (0.1 mM final concentration, C.C.Pro Z-41-M) and 10 mg gentamicin (Sigma G1397) at 4% hematocrit and under 5% O2, 5% CO2 at 37 °C. Parasites were synchronized by sorbitol (5%, Sigma S6021) lysis during ring stage followed by a plasmagel (Plasmion, Fresenius Kabi) enrichment for late blood stages 24 hours later. Another sorbitol treatment 6 h afterwards places the 0 h time point 3 h after the plasmagel enrichment. Parasite development was monitored by Giemsa staining. Parasites were harvested at 1–5% parasitemia.

### Generation of strains

The SMC3-3HA-*glms* strain was generated using a two-plasmid system (pUF1 and pL7) based on the CRISPR/Cas9 system previously described in (Ghorbal et al., 2014). A 3D7 wild-type bulk ring stage culture was transfected with 25 μg pUF1-Cas9 and 25 μg of pL7-*Pf*SMC3-3HA-*glmS* containing the single guide RNA (sgRNA) -encoding sequence 5’- CCTAGAAAATTAGAACAATT-3’ targeting the 3’ UTR of PF3D7_0414000. The pL7-*Pf*SMC3-3HA-*glmS* plasmid also contained the homology repair construct 5’- AGATAGAGAGAGTTATATATCTAAAGGAACAAAGAATGAGGCCTACGAAATTATTAGCATTGTATAAAAAAAAAAAGAAAAAAAAAAGAAAAAAAAAAAAGATTATATATATAATATATGTTGACAATTAATAAATATATTTGTATATATCTGTTAACTAATTATGAAAATTTTTGAATCAATAAATTTTTTAAATAACAAAAAAAAAAAAAAATATATATATTATATATATATTTTATATTTTATATTTTCTTGTAATTTTTGTTTTTTTAGGAGGAAAAACATGCCCTAGAAAATggcggtggaTACCCTTACGATGTGCCTGATTACGCGTAtCCcTAtGAcGTaCCaGAcTAtGCGTACCCtTAtGAcGTtCCgGATTAtGCtcacggggtgTAAGCGGCCGCGGTCTTGTTCTTATTTTCTCAATAGGAAAAGAAGACGGGATTATTGCTTTACCTATAATTATAGCGCCCGAACTAAGCGCCCGGAAAAAGGCTTAGTTGACGAGGATGGAGGTTATCGAATTTTCGGCGGATGCCTCCCGGCTGAGTGTGCAGATCACAGCCGTAAGGATTTCTTCAAACCAAGGGGGTGACTCCTTGAACAAAGAGAAATCACATGATCTTCCAAAAAACATGTAGGAGGGGACAACAATTTGGTTTTGTTTTTTTCTTTAGGTTTTGAGAAAAACAAATAGGAAATACAAAAAAAAAAAAAAAAAAAAAAAAAAAAAAAAAAAAAATGTATTTTTACATATGCACTTGGATTATTTTATTTTTATTATTTTTCTTTATATAAAGTAAAAATATACATAAGTATGCTTATTTATTACATAAGAGTTTATTTAAGAAAGGTTTCTTTTTCATAATATTGTGTGCATGAGTTTTTTTTTATTTTATTTTTTTTTTTTATTTCTGTAACGAAAAGGATATTAAAAAAAATAATAAAA- 3’ (synthesized by GenScript Biotech [Piscataway, NJ, USA]). This homology repair construct comprises a 3 x Hemaglutinin (3HA) - encoding sequence followed by a *glmS* ribozyme-encoding sequence (Prommana et al., 2013), which are flanked by 300 bp homology repair regions upstream and downstream of the Cas9 cut site, excluding the gene STOP codon. The sgRNA sequence was designed using Protospacer (MacPherson & Scherf, 2015). The sgRNA sequence uniquely targeted a single sequence in the genome. As the sgRNA sequence encompasses the STOP codon, its modification via the addition of the 3HA and *glmS*-encoding sequences renders the modified parasites refractory to further dCas9 cleavage at this locus. All cloning was performed using KAPA HiFi DNA Polymerase (Roche 07958846001), In-Fusion HD Cloning Kit (Clontech 639649), and XL10-Gold Ultracompetent E. coli (Agilent Technologies 200315). After transfection, drug selection was applied for five days at 2.67 nM WR99210 (Jacobus Pharmaceuticals) and 1.5 μM DSM1 (MR4/BEI Resources). Parasites reappeared approximately three weeks after transfection, and 5-fluorocytosine was used to negatively select the pL7 plasmid. Parasites were cloned by limiting dilution, and the targeted genomic locus was sequenced to confirm epitope tag and ribozyme integration.

### SMC3 immunoprecipitation and mass spectrometry

An SMC3-3HA-*glms* clone (*n =* 3 technical replicates) and wild-type culture (*n =* 3 technical replicates), as a negative control, were synchronized. Late stage parasites (1.5 × 10^9^ parasites) were enriched using Percoll density gradient separation and then cross-linked with 1 mL 0.5 mM dithiobissuccinimidyl propionate (DSP; Thermo Fisher 22585) in DPBS for 60 min at 37°C (as in (Mesén-Ramírez et al., 2016)). Cross-linked parasites were centrifuged at 4,000 *g* for 5 min at 4°C, and the pellet was washed twice with DPBS at 4°C. The pellet was lysed with 10 volumes of RIPA buffer (10 mM Tris–HCl pH 7.5, 150 mM NaCl, 0.1% SDS, 1% Triton) containing protease and phosphatase inhibitor cocktail (Thermo Fisher 78440) and 1 U/μL of Benzonase (Merck 71206). The lysates were cleared by centrifugation at 16,000 *g* for 10 min at 4°C. Supernatants were incubated with 25 μL of anti-HA Dynabeads (Thermo Fisher 88836) overnight with rotation at 4°C. Beads were collected with a magnet and washed five times with 1 mL RIPA buffer, then five times with 1 mL DPBS, and then once with 1 mL 25 mM NH_4_HCO_3_ (Sigma 09830). The beads were reduced with 100 mM dithiothreitol (Sigma D9779), alkylated with 55 mM iodoacetamide (Sigma I1149), and subjected to on-bead digestion using 1 μg of trypsin (Thermo Fisher 90059). The resulting peptides were desalted using C18 ziptips (Merck ZTC04S096) and sent for MS analysis.

Peptides were separated by reverse phase HPLC (Thermo Fisher Easy-nLC1000) using an EASY-Spray column, 50 cm × 75 μm ID, PepMap RSLC C18, 2 μm (Thermo Fisher ES803A) over a 70-min gradient before nanoelectrospray using a Q Exactive HF-X mass spectrometer (Thermo Fisher). The mass spectrometer was operated in a data-dependent mode. The parameters for the full scan MS were as follows: resolution of 60,000 across 350–1,500 *m*/*z*, AGC 1e^5^ (as in (Kensche et al., 2016)), and maximum injection time (IT) 150 ms. The full MS scan was followed by MS/MS for the top 15 precursor ions in each cycle with an NCE of 30 and dynamic exclusion of 30 s and maximum IT of 96 ms. Raw mass spectral data files (.raw) were searched using Proteome Discoverer 2.3.0.523 (Thermo Fisher) with the SEQUEST search engine. The search parameters were as follows: 10 ppm mass tolerance for precursor ions; 0.8 Da fragment ion mass tolerance; two missed cleavages of trypsin; fixed modification was carbamidomethylation of cysteine; and variable modifications were methionine oxidation, CAMthiopropanoyl on lysine or protein N-terminal, and serine, threonine, and tyrosine phosphorylation. Only peptide spectral matches (PSMs) with an XCorr score greater than or equal to 2 and an isolation interference less than or equal to 30 were included in the data analysis.

### Protein fractionation and western blot analysis

Parasites were washed once with Dulbecco’s phosphate-buffered saline (DPBS, Thermo Fisher 14190), then resuspended in cytoplasmic lysis buffer (25 mM Tris–HCl pH 7.5, 10 mM NaCl, 1.5 mM MgCl_2_, 1% IGEPAL CA-630, and 1× protease inhibitor cocktail [“PI”, Roche 11836170001]) at 4°C and incubated on ice for 30 min. The cytoplasmic lysate was cleared with centrifugation (13,500 *g*, 10 min, 4°C). The pellet (containing the nuclei) was resuspended in 3.3 times less volume of nuclear extraction buffer (25 mM Tris–HCl pH 7.5, 600 mM NaCl, 1.5 mM MgCl_2_, 1% IGEPAL CA-630, PI) than cytoplasmic lysis buffer at 4°C, transferred to 1.5 mL sonication tubes (Diagenode C30010016, 300 µL per tube), and sonicated for five min total (10 cycles of 30 s on/off) in a Diagenode Pico Bioruptor at 4°C. This nuclear lysate was cleared with centrifugation (13,500 *g*, 10 min, 4°C). Protein samples were supplemented with NuPage Sample Buffer (Thermo Fisher NP0008) and NuPage Reducing Agent (Thermo Fisher NP0004) and denatured for 10 min at 70°C. Proteins were separated on a 4–12% Bis-Tris NuPage gel (Thermo Fisher NP0321) and transferred to a PVDF membrane with a Trans-Blot Turbo Transfer system (Bio-Rad). The membrane was blocked for 1 h with 1% milk in PBST (PBS, 0.1% Tween 20) at 25°C. HA-tagged proteins and histone H3 were detected with anti-HA (Abcam ab9110, 1:1,000 in 1% milk-PBST) and anti-H3 (Abcam ab1791, 1:2,500 in 1% milk-PBST) primary antibodies, respectively, followed by donkey anti-rabbit secondary antibody conjugated to horseradish peroxidase (“HRP”, Sigma GENA934, 1:5,000 in 1% milk-PBST). Aldolase was detected with anti-aldolase-HRP (Abcam ab38905, 1:5,000 in 1% milk-PBST). HRP signal was developed with SuperSignal West Pico Plus chemiluminescent substrate (Thermo Fisher 34580) and imaged with a ChemiDoc XRS+ (Bio-Rad).

### Immunofluorescence assays and image acquisition

iRBCs were washed once with DPBS (Thermo Fisher 14190) at 37°C and fixed in suspension in 4% paraformaldehyde (EMS 15714) with 0.0075% glutaraldehyde (EMS 16220) in PBS for 20 min at 25°C, as described previously (Tonkin et al., 2004). The subsequent steps were performed at 25 °C as described in (Mehnert et al., 2019), with minor changes. After washing once with PBS, cells were permeabilized with 0.1% Triton-X 100 for 10 min followed by three PBS washes. Free aldehyde group were quenched with 50 mM NH_4_Cl for 10 min, followed by two PBS washes. Cells were blocked with 3% bovine serum albumin (BSA) (Sigma A4503-50G) in PBS for 30 min. Cells were incubated with anti-HA (Abcam ab9110, 1:1,000 in 3% BSA in PBS) primary antibody for one hour followed by three 10 min washes with 0.5% Tween^®^ 20/PBS. Cells were incubated with anti-rabbit Alexa Fluor 488- or 633-conjugated secondary antibodies (Invitrogen A-11008 or A-21070, 1:2,000 in 3% BSA in PBS) with DAPI (FluoProbes FP-CJF800, 1 μg/mL) for 45 min followed by three 10 min washes with 0.5% Tween^®^ 20/PBS. Cells were washed once more with PBS and placed onto polyethyleneimine-coated slides (Thermo Scientific 30-42H-RED-CE24). Once adhered to the slide, cells were washed twice and mounted with VectaShield® (Vector Laboratories). Images were acquired using a Deltavision Elite imaging system (GE Healthcare), and Fiji (http://fiji.sc) was used for analysis using the least manipulation possible.

### SMC3 chromatin immunoprecipitation sequencing and data analysis

A clonal population of SMC3-3HA-*glmS* parasites were tightly synchronized and harvested at 12 (10^10^ parasites), 24 (4.3 x 10^8^ parasites) and 36 hpi (3.6 x 10^8^ parasites). Parasite culture was centrifuged at 800 *g* for 3 min at 25°C. Medium was removed and the RBCs were lysed with 10 mL 0.075% saponin (Sigma S7900) in DPBS at 37°C. The parasites were centrifuged at 3,250 *g* for 3 min at 25°C and washed with 10 mL DPBS at 37°C. The supernatant was removed, and the parasite pellet was resuspended in 10 mL of PBS at 25°C. The parasites were cross-linked by adding methanol-free formaldehyde (Thermo Fisher 28908) (final concentration 1%) and incubating with gentle agitation for 10 min at 25°C. The cross-linking reaction was quenched by adding glycine (final concentration 125 mM, Sigma G8899) and incubating with gentle agitation for 5 min at 25°C. Parasites were centrifuged at 3,250 *g* for 5 min at 4°C and the supernatant removed. The pellet was washed with DPBS and centrifuged at 3,250 *g* for 5 min at 4°C. The supernatant was removed, and the cross-linked parasite pellet were snap-frozen.

For each time-point, 200 µL of Protein G Dynabeads (Invitrogen 10004D) were washed twice with 1 mL ChIP dilution buffer (16.7 mM Tris–HCl pH 8, 150 mM NaCl, 1.2 mM EDTA pH 8, 1% Triton X-100, 0.01% SDS) using a DynaMag magnet (Thermo Fisher 12321D). The beads were resuspended in 1 mL ChIP dilution buffer with 8 μg of anti-HA antibody (Abcam ab9110) and incubated on a rotator at 4°C for 6 h.

The cross-linked parasites were resuspended in 4 mL of lysis buffer (10 mM HEPES pH 8, 10 mM KCl, 0.1 mM EDTA pH 8, PI) at 4°C, and 10% Nonidet-P40 was added (final concentration 0.25%). The parasites were lysed in a prechilled dounce homogenizer (200 strokes for 12 hpi parasites and 100 strokes for 24 and 36 hpi parasites). The lysates were centrifuged for 10 min at 13,500 *g* at 4°C, the supernatant was removed, and the pellet was resuspended in SDS lysis buffer (50 mM Tris–HCl pH 8, 10 mM EDTA pH 8, 1% SDS, PI) at 4°C (3.6 mL for the 12 hpi sample and 1.8 mL for the 24 ad 36 hpi samples). The liquid was distributed into 1.5 mL sonication tubes (Diagenode C30010016, 300 µL per tube) and sonicated for 12 min total (24 cycles of 30 s on/off) in a Diagenode Pico Bioruptor at 4°C. The sonicated extracts were centrifuged at 13,500 *g* for 10 min at 4°C and the supernatant, corresponding to the chromatin fraction, was kept. The DNA concentration for each time point was determined using the Qubit dsDNA High Sensitivity Assay Kit (Thermo Fisher Scientific Q32851) with a Qubit 3.0 Fluorometer (Thermo Fisher Scientific). For each time point, chromatin lysate corresponding to 100 ng of DNA was diluted in SDS lysis buffer (final volume 200 μL) and kept as “input” at −20°C. Chromatin lysate corresponding to 19 μg (12 hpi), 2 μg (24 hpi) and 3 μg (36 hpi) of DNA was diluted 1:10 in ChIP dilution buffer at 4°C.

Using a DynaMag magnet, the antibody-conjugated Dynabeads were washed twice with 1 mL ChIP dilution buffer and resuspend in 100 μL of ChIP dilution buffer at 4°C. Then the washed antibody-conjugated Dynabeads were added to the diluted chromatin sample and incubated overnight with rotation at 4°C. The beads were collected on a DynaMag into eight different tubes per sample, the supernatant was removed, and the beads in each tube were washed for 5 min with gentle rotation with 1 mL of the following buffers, sequentially:

- Low salt wash buffer (20 mM Tris–HCl pH 8, 150 mM NaCl, 2 mM EDTA pH 8, 1% Triton X-100, 0.1% SDS) at 4°C.
- High salt wash buffer (20 mM Tris–HCl pH 8, 500 mM NaCl, 2 mM EDTA pH 8, 1% Triton X-100, 0.1% SDS) at 4°C.
- LiCl wash buffer (10 mM Tris–HCl pH 8, 250 mM LiCl, 1 mM EDTA pH 8, 0.5% IGEPAL CA-630, 0.5% sodium deoxycholate) at 4°C.
- TE wash buffer (10 mM Tris–HCl pH 8, 1 mM EDTA pH 8) at 25°C.

After the washes, the beads were collected on a DynaMag, the supernatant was removed, and the beads for each time point were resuspended in 800 μL of elution buffer and incubated at 65°C for 30 min with agitation (1000 rpm 30 s on/off). The beads were collected on a DynaMag and the eluate, corresponding to the “ChIP” samples, was transferred to a different tube.

For purification of the DNA, both “ChIP” and “Input” samples were incubated for approximately 10 h at 65°C to reverse the crosslinking. 200 μL of TE buffer followed by 8 μL of RNaseA (Thermo Fisher EN0531) (final concentration of 0.2 mg/mL) were added to each sample, which was then incubated for 2 h at 37 °C. 4 μL Proteinase K (New England Biolabs P8107S) (final concentration of 0.2 mg/mL) were added to each sample, which was then incubated for 2 h at 55°C. 400 μL phenol:chloroform:isoamyl alcohol (25:24:1) (Sigma, 77617) were added to each sample, which was then mixed with vortexing and centrifuged for 10 min at 13,500 *g* at 4°C to separate phases. The aqueous top layer was transferred to another tube and mixed with 30 μg glycogen (Thermo Fisher 10814) and 5M NaCl (200 mM final concentration). 800 μL 100% EtOH at 4°C were added to each sample, which was then incubated at −20°C for 30 min. The DNA was pelleted by centrifugation for 10 min at 13,500 *g* at 4°C, washed with 500 μL 80% EtOH at 4°C, and centrifuged for 5 min at 13,500 *g* at 4°C. After removing the EtOH, the pellet was dried at 25 °C and all DNA for each sample was resuspended in 30 μL 10 mM Tris–HCl, pH 8 total. The DNA concentration and average size of the sonicated fragments was determined using a DNA high sensitivity kit and the Agilent 2100 Bioanalyzer. Libraries for Illumina Next Generation Sequencing were prepared with the MicroPlex library preparation kit (Diagenode C05010014), with KAPA HiFi polymerase (KAPA biosystems) substituted for the PCR amplification. Libraries were sequenced on the NextSeq 500 platform (Illumina).

Sequenced reads (150 bp paired end) were mapped to the *P. falciparum* genome (Gardner et al., 2002) (plasmoDB.org, version 3, release 55) using “bwa mem” (Li & Durbin, 2009) allowing a read to align only once to the reference genome (option “–c 1”). Alignments were subsequently filtered for duplicates and a mapping quality ≥ 20 using samtools (Li et al., 2009). The paired end deduplicated ChIP and input BAM files were used as treatment and control, respectively, for peak calling with the macs2 command callpeak default settings (Y. Zhang et al., 2008). Obtained peaks with *q*-value cutoff 0.05 for each time point were visualized in Integrative Genomics Viewer (Robinson et al., 2011) along with ChIP-Input ratio coverage obtained from deeptool’s bamCompare command (Ramírez et al., 2016). To map SMC3 binding to nearby protein coding genes, peak summits were extended 150 bp upstream and downstream, and bedtools closest command (Quinlan & Hall, 2010) were used with *P. falciparum* reference genome feature file (gff) (plasmoDB.org, version 3, release 56). Only regions 500 bp upstream or downstream near to the protein coding genes were considered further for downstream analysis. Centromeric regions (from (Hoeijmakers et al., 2012) were corrected for changes in genome annotation. These regions were overlapped with SMC3 peaks dataset using bedtools intersect command (Quinlan & Hall, 2010). Fold change quantification and statistical analysis for all peaks and peaks in centromeric regions was performed in R (R Core Team, 2021).

### RNA extraction, stranded RNA sequencing and analysis

A WT and SMC3-3HA-*glmS* clone were synchronized simultaneously and each culture was split into two at 12 hpi. Glucosamine (Sigma G1514, final concentration 2.5 nM) was added to one half of the culture for two rounds of parasite replication (approximately 96 h). Parasites were then re-synchronized and three technical replicates (with and without glucosamine) were harvested at 12, 24, and 36 hpi. RBCs were lysed in 0.075 % saponin (Sigma S7900) in PBS at 37°C, centrifuged at 3,250 *g* for 5 min, washed in PBS, centrifuged at 3,250 g for 5 min, and resuspended in 700 μL QIAzol reagent (Qiagen 79306). RNA was extracted using an miRNeasy Mini kit (Qiagen 1038703) with the recommended on-column DNase treatment. Total RNA was poly (A) selected using the Dynabeads mRNA Purification Kit (Thermo Fischer Scientific 61006). Library preparation was performed with the NEBNext® Ultra™ II Directional RNA Library Prep Kit for Illumina® (New England Biolabs E7760S) and paired end sequencing was performed on the Nextseq 550 platform (Illumina). Sequenced reads (150 bp paired end) were mapped to the *P. falciparum* genome (Gardner et al., 2002) (plasmoDB.org, version 3, release 55) using “bwa mem” (Li & Durbin, 2009), allowing a read to align only once to the reference genome (option “–c 1”). Alignments were subsequently filtered for duplicates and a mapping quality ≥ 20 using samtools (Li et al., 2009). Gene counts were quantified with htseq-count (Anders et al., 2015), and differentially expressed genes were identified in R (R Core Team, 2021) using package DESeq2 (Love et al., 2014).

### Estimation of cell cycle progression

RNA-seq-based cell cycle progression was estimated in R by comparing the normalized expression values (i.e., RPKM, reads per kilobase per exon per one million mapped reads) of each sample to the microarray data from (Bozdech et al., 2003) using the statistical model as in (Lemieux et al., 2009).

### Parasite growth assay

Parasite growth was measured as described previously (Vembar et al., 2015). Briefly, an SMC3-3HA-*glmS* clone and a WT clone were tightly synchronized. Each culture was split and glucosamine (Sigma G1514) was added (2.5 mM final concentration) to one half for approximately 96 h before starting the growth curve. The parasites were tightly re-synchronized and diluted to 0.3% parasitemia (5% hematocrit) at ring stage using the blood of two different donors. The growth curve was performed in a 96-well plate (200 μL culture per well) with three technical replicates per condition per blood. Every 24 h, 5 μL of the culture were fixed in 45 μL of 0.025% glutaraldehyde in PBS for 1h at 4°C. After centrifuging at 800 *g* for 5 min, free aldehyde groups were quenched by re-suspending the iRBC pellet in 200 μL of 15 mM NH_4_Cl in PBS. A 1:10 dilution of the quenched iRBC suspension was incubated with Sybr Green I (Sigma S9430) to stain the parasite nuclei. Quantification of the iRBCs was performed in a CytoFLEX S cytometer (Beckman Coulter) and analysis with FlowJo™ Software.

## DATA AVAILABILITY

All data sets generated in this study are available in the following databases:

- ChIP-seq data: NCBI BioProject accession # PRJNA854331
- RNA-seq data: NCBI BioProject accession # PRJNA854331
- SMC3-3HA Proteomics data: PRIDE repository accession # PXD035225 (DOI: 10.6019/PXD035225, Username, Password: agXjJYP4)

## ACKNOWLEDGEMENTS

This work was supported by the Agence Nationale de la Recherche (grant ANR-21-CE15-0010 PlasmoVarOrg to JMB and grant ANR-11-LABEX-0024-01 ParaFrap to ASc); the European Research Council (grant PlasmoSilencing 670301 to ASc and grant PlasmoEpiRNA 947819 to SB); the Academic Research Fund (Tier 2) of the Ministry of Education, Singapore (MOE2018-T2-2-131 to PRP); and the Merlion Project (6.11.18 to PRP). Work in the laboratories of PRP and PCD was funded by the National Research Foundation of Singapore through the Singapore-MIT Alliance for Research and Technology Antimicrobial Resistance Interdisciplinary Research Group. JMB was supported by a European Molecular Biology Organization long-term postdoctoral fellowship (EMBO ALTF 180-2015), the Institut Pasteur Roux-Cantarini postdoctoral fellowship, and a ParaFrap fellowship. SB was supported by a European Molecular Biology Organization long-term postdoctoral fellowship (EMBO ALTF 1444-2016) and advanced fellowship (EMBO aALTF 632-2018). ASi was supported by the Singapore-MIT Alliance (SMA) Graduate Fellowship from the Ministry of Education of Singapore. The authors would like to acknowledge the use of the Biomics and Flow Cytometry platforms at the Institut Pasteur, as well as the mass spectrometer facilities at A*STAR Institute of Molecular and Cell Biology in the laboratory of Dr. Radoslaw Sabota with the aid of Dr. Wint Wint Phoo. The authors would like to thank Aurélie Claës and Patty Chen of the Biology of Host Parasite Interactions unit at the Institut Pasteur for their invaluable support in the lab.

## AUTHOR CONTRIBUTIONS

CR and JMB conceptualized the project and conceived experiments. PS and SB performed DNA/RNA sequencing bioinformatic analysis. ASi performed the SMC3 immunoprecipitation mass spectrometry and analysis. CR performed all other experiments. PRP, PCD, and ASc supervised and helped interpret analyses. CR, PS, and JMB wrote the manuscript. All authors discussed and approved the manuscript.

## TABLE LEGENDS

**Table 1. LC-MS/MS analysis of SMC3-3HA immunoprecipitation.** LC-MS/MS results of the SMC3-3HA immunoprecipitation in late stage parasites. Total (TotPep) and unique (UniPep) peptide counts for the proteins listed are shown for three replicates each of the SMC3-3HA and 3D7 WT control immunoprecipitations. Predicted members of the cohesin complex are highlighted in grey based on (Hillier et al., 2019; Batugedara et al., 2020).

**Table 2. MACS2 peak calling results for SMC3-3HA ChIP-seq at 12, 24, and 36 hpi.** The paired end deduplicated ChIP and input BAM files were used as treatment and control, respectively, for peak calling algorithm macs2 command callpeak. Significant peaks (*q* < 0.05) are shown for each time point, along with their chromosomal coordinates, fold enrichment (ChIP/Input), and -log_10_(*q*-value).

**Table 3. SMC3-3HA peak enrichment at centromeric regions at 12, 24, and 36 hpi.** List of significant SMC3 peaks (*q* < 0.05, Table 2) that overlap with centromeres, as defined by peaks of CenH3 (Hoeijmakers et al., 2012) at 12, 24, and 36 hpi. Significant SMC3 peaks and their overlapping centromeric regions are shown for each time point, along with their chromosomal coordinates, fold enrichment (ChIP/Input), and -log_10_(*q*-value).

**Table 4. List of SMC3-3HA peak-associated genes at 12, 24, and 36 hpi.** Protein-coding genes that are closest to the SMC3-3HA peak summit (+/- 500 bp) at 12, 24, and 36 hpi (defined in Table 2).

**Table 5. Gene Ontology analysis for SMC3 peak-associated genes.** GO enrichment analysis (biological process) of genes associated with an SMC3 peak at 12, 24, or 36 hpi (defined in Table 4). Number of significantly enriched genes within each “biological process” term (Result count), number of genes with this term divided by the total number of annotated genes with this term in the *P. falciparum* genome (Fold enrichment), odds ratio statistics from the Fisher’s exact test, *P*-value (calculated using a Fisher’s exact test), and Benjamini-corrected *P*-value are shown (*q*-value). Only GO terms with *P* < 0.05 are shown. Analysis was performed using the GO enrichment tool at PlasmoDB.org (Aurrecoechea et al., 2017).

**Table 6. Differential gene expression analysis at 12 hpi of glucosamine-treated over untreated SMC3-3HA-*glmS* parasites.** Analysis was performed for n=3153 genes (ID and chromosome locations are given) with two and three replicates for untreated and glucosamine-treated SMC3-3HA-*glmS* parasites, respectively. SMC3 is highlighted in grey. log_2_(FoldChange) = Fold change of baseMean (average of the normalized read counts across all samples and replicates for this gene) in glucosamine-treated/untreated parasites (log_2_). *P*-values are calculated with a Wald test for significance of coefficients in a negative binomial generalized linear model as implemented in DESeq2 (Love et al., 2014). *q* = Bonferroni corrected *P-*value.

**Table 7. Differential gene expression analysis at 24 hpi of glucosamine-treated over untreated SMC3-3HA-*glmS* parasites.** Analysis was performed for n=4822 genes (ID and chromosome locations are given) with three replicates for untreated and glucosamine-treated SMC3-3HA-*glmS* parasites. SMC3 is highlighted in grey. log_2_(FoldChange) = Fold change of baseMean (average of the normalized read counts across all samples and replicates for this gene) in glucosamine-treated/untreated parasites (log_2_). *P*-values are calculated with a Wald test for significance of coefficients in a negative binomial generalized linear model as implemented in DESeq2 (Love et al., 2014) *q* = Bonferroni corrected *P*-value.

**Table 8. Differential gene expression analysis at 12 hpi of glucosamine-treated over untreated 3D7 WT parasites.** Analysis was performed for n=3668 genes (ID and chromosome locations are given) with three replicates for untreated and glucosamine-treated 3D7 WT parasites, respectively. log_2_(FoldChange) = Fold change of baseMean (average of the normalized read counts across all samples and replicates for this gene) in glucosamine-treated/untreated parasites (log_2_). *P*-values are calculated with a Wald test for significance of coefficients in a negative binomial generalized linear model as implemented in DESeq2 (Love et al., 2014). *q* = Bonferroni corrected *P*-value.

**Table 9. Differential gene expression analysis at 24 hpi of glucosamine-treated over untreated 3D7 WT parasites.** Analysis was performed for n=4734 genes (ID and chromosome locations are given) with three replicates for untreated and glucosamine-treated 3D7 WT parasites. log_2_(FoldChange) = Fold change of baseMean (average of the normalized read counts across all samples and replicates for this gene) in glucosamine-treated/untreated parasites (log_2_). *P*-values are calculated with a Wald test for significance of coefficients in a negative binomial generalized linear model as implemented in DESeq2 (Love et al., 2014). *q* = Bonferroni corrected *P*-value.

**Table 10. List of differentially expressed genes in SMC3-3HA-*glmS* parasites after filtering of significantly differentially expressed genes in the 3D7 WT upon glucosamine treatment at 12 hpi (Supp. Fig. 2).** log_2_(FoldChange) = Fold change of baseMean (average of the normalized read counts across all samples and replicates for this gene) in glucosamine-treated/untreated parasites (log_2_). *P*-values are calculated with a Wald test for significance of coefficients in a negative binomial generalized linear model as implemented in DESeq2 (Love et al., 2014). *q* = Bonferroni corrected *P*-value. SMC3 is highlighted in grey.

**Table 11. List of differentially expressed genes in SMC3-3HA-*glmS* parasites after filtering of significantly differentially expressed genes in the 3D7 WT upon glucosamine treatment at 24hpi (Supp. Fig. 2).** log_2_(FoldChange) = Fold change of baseMean (average of the normalized read counts across all samples and replicates for this gene) in glucosamine-treated/untreated parasites (log_2_). *P*-values are calculated with a Wald test for significance of coefficients in a negative binomial generalized linear model as implemented in DESeq2 (Love et al, 2014). *q* = Bonferroni corrected *P*-value. SMC3 is highlighted in grey.

**Table 12: Gene Ontology analysis of significantly up- and downregulated genes in SMC3 knockdown at 12 hpi.** GO enrichment analysis (biological process) of genes significantly and specifically up- or downregulated upon SMC3 knockdown at 12 hpi (as defined in Table 10). Number of significantly enriched genes within each “biological process” term (Result count), number of genes with this term divided by the total number of annotated genes with this term in the *P. falciparum* genome (Fold enrichment), odds ratio statistics from the Fisher’s exact test, *P*-value (calculated using a Fisher’s exact test), and Benjamini-corrected *P*-value (*q*-value). Only GO terms with *P* < 0.05 are shown. Analysis was performed using the GO enrichment tool at PlasmoDB.org (Aurrecoechea et al., 2017).

**Table 13: Gene Ontology analysis of significantly up- and downregulated genes in SMC3 knockdown at 24 hpi.** GO enrichment analysis (biological process) of genes significantly and specifically up- or downregulated upon SMC3 knockdown at 12 hpi (as defined in Table 11). Number of significantly enriched genes within each “biological process” term (Result count), number of genes with this term divided by the total number of annotated genes with this term in the *P. falciparum* genome (Fold enrichment), odds ratio statistics from the Fisher’s exact test, *P*-value (calculated using a Fisher’s exact test), and Benjamini-corrected *P*-value (*q*-value). Only GO terms with a *P* < 0.05 are shown. Analysis was performed using the GO enrichment tool at PlasmoDB.org (Aurrecoechea et al., 2017).

**Table 14: List of invasion-related genes that are significantly upregulated in SMC3 knockdown at 12 and 24 hpi.** List of genes significantly and specifically upregulated at 12 and 24 hpi in response to SMC3 depletion that overlap with a list of “invasion-related genes,” as defined in (Hu et al., 2010). Gene IDs and chromosome locations are given. log_2_(FoldChange) = Fold change of baseMean (average of the normalized read counts across all samples and replicates for this gene) in glucosamine-treated/untreated parasites (log_2_). *P*-values are calculated with a Wald test for significance of coefficients in a negative binomial generalized linear model as implemented in DESeq2 (Love et al., 2014). *q* = Bonferroni corrected *P*-value.

## SUPPLEMENTARY FIGURE LEGENDS

**Supplementary Figure 1. SMC3 knockdown occurs after two cell cycles of glucosamine treatment.**

Western blot analysis of nuclear extracts from a synchronous clonal population of SMC3-3HA-*glmS* ring stage parasites in the absence (-) or presence (+) of glucosamine (GlcN) for 48 and 96 hr (one and two IDC cycles, respectively). SMC3-3HA is detected with an anti-HA antibody. An antibody against histone H3 is used as a control for the nuclear extract. Molecular weights are shown to the right.

**Supplementary Figure 2. Strategy for determining expression changes due to glucosamine treatment.**

Venn diagram showing the number of unique or shared significantly up- or downregulated genes after two cycles of glucosamine treatment in synchronous, clonal populations of SMC3-3HA-*glmS* (green) and WT (purple) parasites at 12 and 24 hpi.

**Supplementary Figure 3. Cell cycle progression of SMC3-3HA-*glmS* at 12 and 24 hpi.**

Cell cycle progression estimation of a synchronous, clonal SMC3-3HA-*glmS* population in the absence (− GlcN) or presence (+ GlcN) of glucosamine. RNA-seq data from synchronized parasites harvested at 12 (blue) and 24 (coral) hpi were compared to microarray data from (Bozdech et al., 2003) as in (Lemieux et al., 2009) to determine the approximate time point in the IDC (*x*-axis). Replicates are represented with circles.

